# Thyroid hormone-dependent apoptosis during metamorphosis in *Ciona robusta* involves both bilaterian-ancestral and vertebrate-derived processes

**DOI:** 10.1101/2023.01.06.522990

**Authors:** Godefroy Nelly, Le Goff Emilie, Pan Qiaowei, Baghdiguian Stephen, Debiais-Thibaud Mélanie, Martinand-Mari Camille

## Abstract

Chordate metamorphosis is a postembryonic larva-to-juvenile transition triggered by thyroid hormones and their specific receptors (TR). This crucial developmental event shows a wide morphological diversity among different chordate lineages and is characterized by ecological, morphological, metabolic and behavioral changes that can be drastic. One of the most studied models is the amphibian Xenopus, whose tadpole metamorphosis includes apoptosis-induced tail regression dependent on the thyroid hormone pathway. In an evolutionary context, we used the ascidian model, the extant closest group to vertebrates, in which the swimming larva transforms to a sessile filter-feeding juvenile during metamorphosis, to study the role of thyroid hormones in this transformation. The ascidian metamorphosis is also characterized by an apoptosis-driven tail regression as in Xenopus. However, whether this apoptosis-driven process is dependent on the thyroid hormone has not yet been elucidated.

In this study, we interfered with thyroid hormone signaling during tail regression of the ascidian *Ciona robusta* to investigate whether (i) thyroid hormone is involved in the regulation of developmental apoptosis, and (ii) apoptosis leading to tail regression involves its classical molecular pathways. We described specific gene expression landmarks as well as apoptosis dynamics during larva metamorphosis under thyroid hormone exposure and thyroid hormone inhibition treatments. We provide evidence that *Ciona robusta* metamorphosis involves thyroid hormone-dependent apoptosis, similar to other studied chordates. However, the mode of action of thyroid hormone shows great variation compared to the classically described scheme in chordates, both in thyroid hormone/TR interactions and in the apoptotic pathway.

## 1. Introduction

The thyroid hormones thyroxin (T4) and its derivative tri-iodothyronine (T3) are essential to organogenesis and tissue rearrangements during metamorphosis, as well as to growth, development, tissue homeostasis and metabolism during the full life cycle of vertebrates (Mullur *et al*., 2014; Seoane-Collazo *et al*., 2015; Goemann *et al*., 2017). Thyroid hormones circulate in the plasma (Roche *et al*., 1962; Barrington & Thorpe, 1965; Fujita & Sawano, 1979; Dunn, 1980a; b; Di Fiore *et al*., 1997) and are activated/catalyzed by deiodinases in the tissues (Darras and Van Herck, 2012). Amphibians have been used as a model to study vertebrate metamorphosis for almost a century. Metamorphosis involves the remodeling of several organ systems which necessitates the orchestration of both cell proliferation and cell death: in this regard, one spectacular event in amphibian metamorphosis is tail regression that involves apoptosis in the muscular, skeletal and nervous systems, regulated by thyroid hormone signaling (Sachs *et al*., 2000a; Shi & Ishizuya-Oka, 2001; Tata, 2006; Ishizuya-Oka, 2011). The molecular mechanisms involved in vertebrate metamorphosis were best described in the studies of amphibians and some teleost fishes. In both models, thyroid hormones function by binding to their nuclear receptors (thyroid hormone receptors, TRα and TRβ) that act as transcription factors to regulate developmental gene expression, including apoptotic genes (for reviews, see Ishizuya-Oka *et al*., 2010; Grimaldi *et al*., 2013; Holzer & Laudet, 2013; Morvan-Dubois *et al*., 2013; Wrutniak-Cabello *et al*., 2017). TR does not usually bind DNA alone but first forms heterodimers with retinoid X receptor (RXRs: RXRα, RXRβ and RXRγ) and then TR/RXR acts as the functional unit to bind DNA and control gene transcription (Ikeda *et al*., 1994; Machado *et al*., 2009). During *Xenopus* tail regression, the thyroid hormone signaling pathway up-regulates *TRβ* expression and down-regulates *RXRγ* expression (Ikeda *et al*., 1994; Machado *et al*., 2009).

Despite a quite complete description of cellular events linked to metamorphosis in Xenopus, including the relationship between TH signaling regulating apoptosis wave(s), there has been little data suggesting this observation is generalized to all vertebrates, or may even be shared with more distant groups. Ascidians represent the extant sister-group to vertebrates and are very easily manipulable in laboratory. Metamorphosis of the ascidian *Ciona robusta* (Brunetti *et al*., 2015) represents a period of profound morphological changes during which the animal alters its life traits (Sasakura & Hozumi, 2017). The lecithotrophic larva has a prototypical chordate body plan, it swims a few hours after hatching and before adhering to the substrate, and then needs a few more hours to metamorphose into a sessile filter-feeding juvenile in laboratory conditions (Matsunobu & Sasakura, 2015). The initiation of metamorphosis is sensitive to several internal and external cues and is morphologically first characterized by tail regression (Matsunobu & Sasakura, 2015), which involves apoptosis (Chambon *et al*., 2002; Tarallo & Sordino, 2004) and cell migration to the trunk region (For review, see Karaiskou *et al*., 2015). To date, whether thyroid hormone plays a conserved role in the regulation of this apoptotic event in ascidians is not known.

In *Ciona robusta* adults, the synthesis of thyroid hormones was suggested to take place in the endostyle, an organ homologous to the thyroid gland of vertebrates (Ogasawara *et al*., 1999). While no endostyle nor blood circulatory system has yet developed in the larvae of *Ciona robusta*, T4 is already present in the tail of 24-hour-old larvae (Patricolo *et al*., 2001). In previous studies, *Ciona robusta* larvae were treated with exogenous T4 or thiourea (TU, an inhibitor of thyroid hormone synthesis), which showed that thyroid hormone is involved in the regulation of metamorphosis (Patricolo *et al*., 2001; D’Agati & Cammarata, 2006). In the genome of *Ciona robusta*, a single TR ortholog was identified. This TR gene contains a highly conserved DNA binding domain, similar to that of vertebrates. The expression of this TR has been detected in the embryo and larva, suggesting a possible role during embryonic development and metamorphosis in *Ciona robusta* (Carosa *et al*., 1998). However, since this receptor was shown to not bind any iodinated tyrosine derivative *in vitro*, the molecular pathways triggered by thyroid hormones to regulated metamorphosis remain elusive (Carosa *et al*., 1998; Paris *et al*., 2008). In the genome of *Ciona robusta*, a single RXR ortholog was identified (Yagi *et al*., 2003). In this regard, *Ciona robusta* is more similar to other deuterostomes or bilaterians (Howard-Ashby *et al*., 2006; Huang *et al*., 2015; Taylor & Heyland, 2018; Li *et al*., 2020).

In this study, we interfered with thyroid hormone signaling during tail regression to characterize whether (i) thyroid hormones are involved in the regulation of apoptosis and (ii) the process of apoptosis involves actors of the classical apoptotic pathways. In order to address these questions, we describe specific gene expression patterns as well as data on apoptosis dynamics during larva metamorphosis under T4 and TU treatments. We provide evidence that thyroid hormone-dependent apoptosis is involved in *Ciona robusta* metamorphosis, similar to previous findings in amphibians. However, the mode of action of thyroid hormone shows great variation compared to the classically described scheme in chordates, both in thyroid hormone/TR interactions and in the apoptotic pathway.

## 2. Material and methods

### 2.1. Ethical statement

The research described herein was performed on *Ciona robusta*, a marine invertebrate. The study was carried out in strict accordance with European and French legislations (directives 2010/63 and 2016-XIX-120, respectively) for the care and use of animals for scientific purposes (ISEM agreement N°A34-172-042) although *Ciona robusta* is not included in the organisms designated by this legislation. The study did not involve endangered nor protected species.

### 2.2. Animal husbandry

Adult *Ciona robusta* were collected in the Thau laguna (SMEL of Montpellier University, France) and maintained at 18°C in a tank with circulating seawater and under constant light to allow gametes accumulation. Oocytes were collected from each individual into separate wells and were fertilized with a mixture of sperms obtained from the different individuals after dissection of gonoducts and spermiducts. Just before fertilization, 50 μl of sperm was activated with 1 ml of Tris 50mM in seawater. Three drops of diluted sperm were used to fertilize each pool of oocytes. Fertilized eggs were reared in tissue culture dishes at 18°C in 0,2 mm filtered seawater containing 100 U/ml penicillin, and 0,1 mg/ml streptomycin (30ml/dish). Embryos and larvae were allowed to develop to the desired stage and then collected or used for further experiments.

### 2.3. Thyroxin, thiourea and caspase inhibitor treatments

Thiourea powder (TU; 16217, Riedel-de Haën) was dissolved directly in seawater to obtain a 500μM working solution. A stock solution of L-thyroxine (T4; T1775, Sigma) was prepared at 1mM in 0,001N NaOH, which was then diluted with seawater to obtain a 100nM working solution.

Just after hatching, swimming larvae (around 100 larvae/kinetic point) were transferred in new tissue culture dishes containing seawater, T4 or TU solutions (30ml/dish) and collected every two hours for 10 hours for qPCR experiments or at 6 and 12h of treatment for rescue experiments (TUNEL staining).

For the tail amputation experiments, the just hatched larvae were anesthetized with MS222 (Sigma-Aldrich, used at 0,8mM final) and 25% of the posterior end of the tails was removed under the binocular with a needle. Afterwards the amputated larvae (50 larvae/kinetic point/treatment) were transferred to tissue culture dishes containing seawater (control) or T4 (100nM in seawater) (30ml/dish). At 6 and 12 hours post treatment, state of larvae was evaluated: no-tail larvae *versus* swimming larvae. The resulting graphs are presented in percentage, 100% corresponding to the total of larvae/plate. Then, larvae were collected and fixed for apoptosis detection by TUNEL staining.

Caspase-8 inhibitor Z-IETD-FMK and Caspase-9 inhibitor Z-LEHD-FMK were obtained from R&D Systems and were dissolved in dimethyl sulfoxide (DMSO, Sigma) to give a 20mM stock solution. Then they were diluted with seawater to obtain a 100μM working solution. At hatching, swimming larvae (100 larvae/condition) were transferred into 24 well plate (500 μl solution/well) and reared in DMSO 0,5% in seawater (DMSO used as control) or caspase inhibitor solutions. The number of tail-regressed larvae was counted 10h after treatment.

### 2.4. TUNEL staining and indirect immunofluorescence analysis

Larvae were fixed for 20 min with 3,7% formaldehyde in filtered seawater and then permeabilized for 20 min at room temperature with 0,2 % Triton X-100 in TS solution (150 mM NaCl, 25 mM Tris, pH 7.5). TUNEL staining (Roche, *In situ* cell death detection kit, TMR red) was performed according to the manufacturer’s instructions. In brief, larvae were incubated in TUNEL reaction mixture (Enzyme solution 10% in Label solution) 1 hour at 37°C in a humidified chamber. Both negative and positive controls of TUNEL staining were performed according to the manufacturer’s instructions. For indirect immunofluorescence, T4 thyroid hormone was detected with a rabbit anti-L-thyroxine polyclonal antibody (T-2652, Sigma), and DNA with DAPI (D9542, Sigma). Appropriate secondary antibody was FITC-conjugated donkey-anti-rabbit immunoglobulins (Jackson Laboratories). Specimens were analyzed with a Leica TCS-SPE laser confocal microscope (Montpellier RIO Imaging platform, France).

### 2.5. RNA isolation, semi-quantitative RT-PCR and qPCR analysis

For semi-quantitative PCR experiment, fertilized eggs were allowed to develop to the desired stage, collected every 2 hours from fertilization to 28 hours post-fertilization (hpf) (100 individuals/kinetic point) and frozen before RNA extraction.

For qPCR experiments, hatched larvae were treated or not with TU and collected every 2 hours to 10 hours post-hatching (hph). At the collecting time, and to focus the study on tail regression, only tails of larvae were collected with a needle under the binocular and frozen before RNA extraction. At each time point, a pool of 100 tails was collected. Total RNA was isolated with RNeasy kit according to the supplier’s instructions (QIAGEN). 70 ng of total RNA were used for cDNA preparation performed by Superscript II reverse transcriptase (Invitrogen) with an oligodT primer, the mixture was incubated for 50 min at 42°C followed by 15 min at 70°C.

Semi-quantitative PCR was performed on cDNA from each time point of kinetic (95°C for 5 min and then 35 cycles of 95°C for 30s, 53°C for 30s, 72°C for 1 min, completed at 72°C for 10 min) and PCR products were run on 2% agarose gels at 100 Volts during 30 min. PCR products were quantified by Image J gel analysis. Each lane was normalized with the expression of reference gene S26 (Vincent *et al*., 1993).

For quantitative PCR, 1:20 dilution of each cDNA was run in triplicate on a 384-well plate for each primer pair by using thermal cycling parameters: 95°C for 10 min, 95°C for 10s, 63°C for 10s, 72°C for 10s (45 cycles) and an additional step 72°C for 10 min performed on a Light Cycler 480 with the SYBR Green I Master kit (Roche) (qPHD UM2/GenomiX Platform, Montpellier - France). Results were normalized with the expression of reference gene S26. Data were analyzed with the Light Cycler 480 software 1.5.1.

All the sequences used come from the Aniseed website (www.aniseed.cnrs.fr). We used Primer 3.0 to design all the sets of forward and reverse primers to amplify selected genes listed in table 1.

**Table 1:**
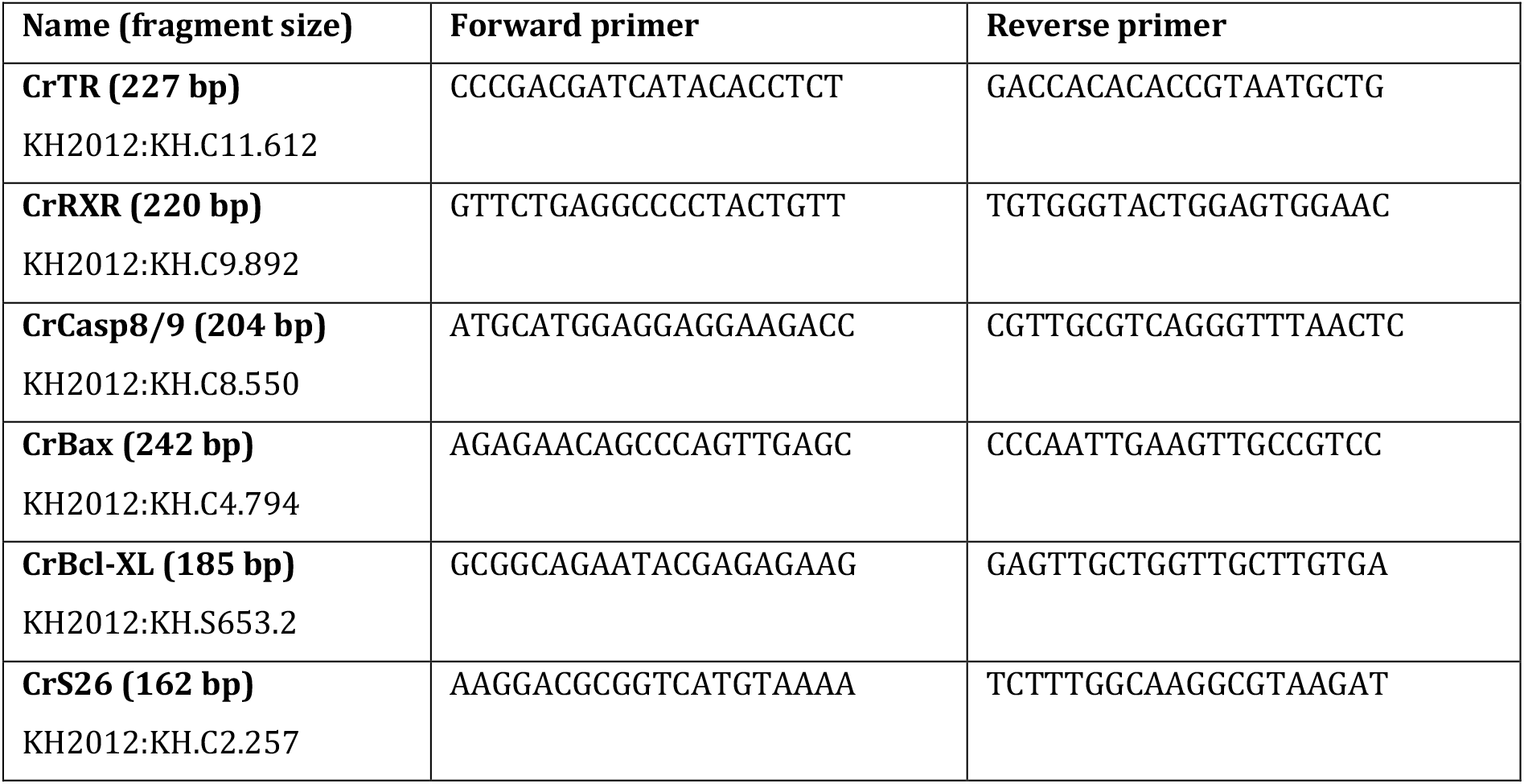
Forward and reverse primers used for qPCR experiments.

### 2.6. Statistical analysis

The semi-quantitative PCR and qPCR experiments were repeated three times (three different spawnings) and the inhibitor treatments were repeated twice. The values are the means +/- standard deviation. At each time point, a Student’s t-test was performed to validate a difference of expression between control and treatment (significant at *P*<0,05). ANOVA was done (for qPCR only) with expression as the response, and time, treatment and their interaction as predictors (Table 2).

**Table 2:** Analysis of Variance (ANOVA) for qPCR experiments. Bold numbers mean *P* value < 0.05.

| gene | Predictor | Sum Sq | Mean Sq | F value | Pr(>F) |
| --- | --- | --- | --- | --- | --- |
| **CrCasp8/9** | time | 0.5663 | 0.5663 | 1.5028 | 0.229180 |
|  | treatment | 4.4219 | 4.4219 | 11.7347 | **0.001702** |
|  | time:treatment | 0.5527 | 0.5527 | 1.4668 | 0.234726 |
| **CrBax** | time | 0.9929 | 0.99294 | 4.0365 | 0.05302 |
|  | treatment | 0.2244 | 0.22439 | 0.9122 | 0.34669 |
|  | time:treatment | 0.0580 | 0.05801 | 0.2358 | 0.63055 |
| **CrBclXI** | time | 53.444 | 53.444 | 39.1998 | **5.11e-07** |
|  | treatment | 3.867 | 3.867 | 2.8363 | 0.1019 |
|  | time:treatment | 3.478 | 3.478 | 2.5509 | 0.1201 |
| **CrRXR** | time | 17.797 | 17.7974 | 16.1577 | **0.0003314** |
|  | treatment | 5.245 | 5.2449 | 4.7617 | **0.0365539** |
|  | time:treatment | 0.053 | 0.0527 | 0.0479 | 0.8282364 |
| **CrTR** | time | 1612.47 | 1612.47 | 87.8360 | **8.037e-10** |
|  | treatment | 372.28 | 372.28 | 20.2793 | **0.0001247** |
|  | time:treatment | 0.25 | 0.25 | 0.0138 | 0.9074141 |

## 3. Results

### 3.1. *An apoptotic wave in tail cells starts after T4 and* CrTR *expression decreases*

The presence of endogenous T4 in the tail of *Ciona robusta* during metamorphosis is detected by immunofluorescence in larvae from hatching (18hpf) to metamorphosis (30hpf). T4 is detectable in all cell-types of the entire tail at hatching, and then gradually disappears from the posterior to the anterior pole of the tail (Figure 1A and supplementary data). No endostyle nor blood circulatory system has yet developed in the larvae and T4 appears diffuse in all tissues of the larva. A massive apoptotic wave is detected through TUNEL staining and first occurs in cells at the posterior extremity of the tail. Then, this death progresses towards the anterior pole together with T4 disappearance, while tail regression has not started yet (25hpf, Figure 1A; see previous results from (Chambon *et al*., 2002). The expression of the single thyroid-hormone receptor *CrTR* is undetectable before 8hpf. Then it increases to reach its maximum at around hatching (Figure 1B, C), and finally decreases until 28hpf.

**Figure 1:**
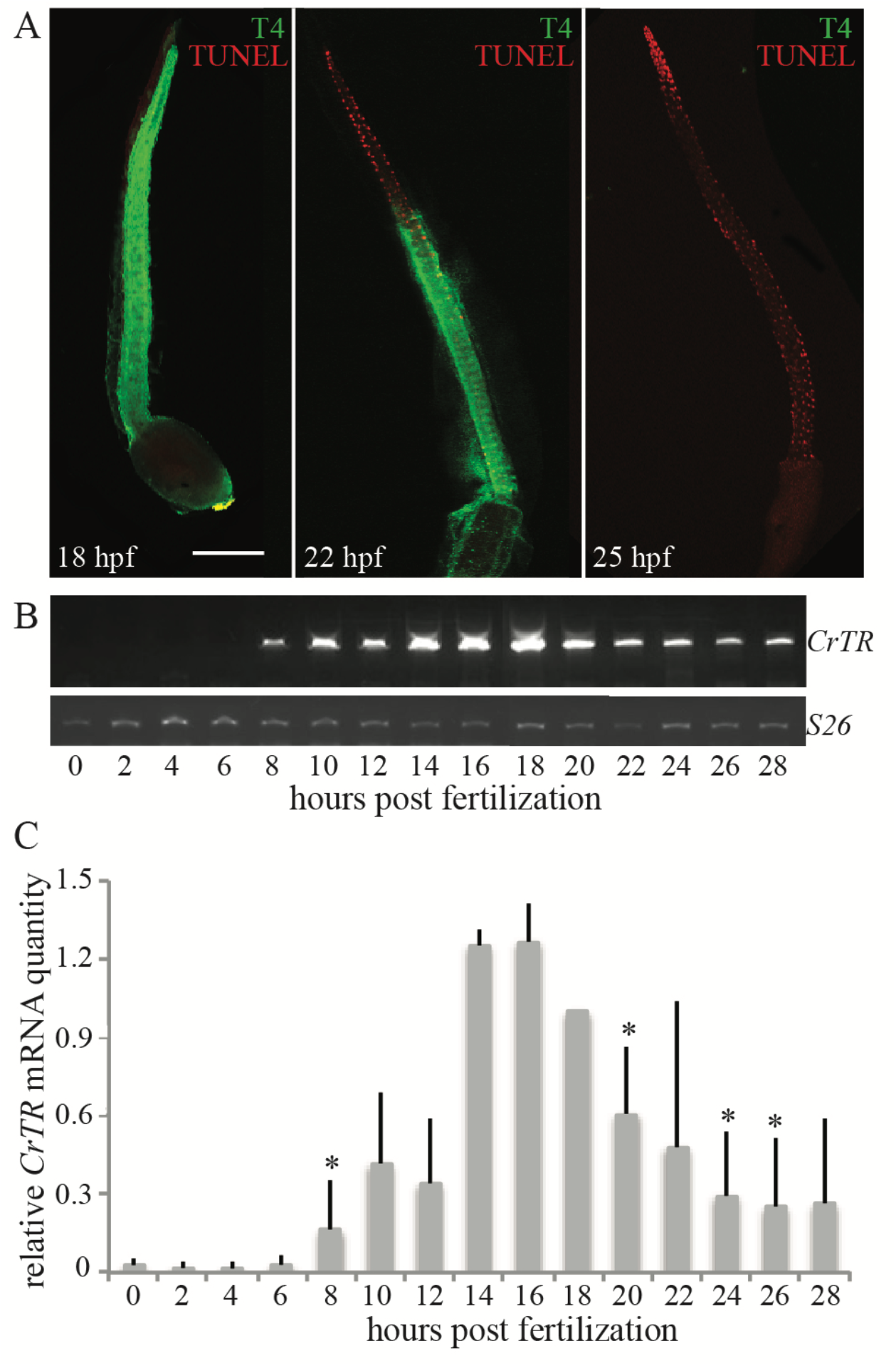
T4 disappearance, *CrTR* expression decrease and apoptosis wave in the tail of *Ciona robusta*. A: TUNEL detection (red nuclei) and T4 immunostaining (green) at different stages of development (hph: hour post-hatching) by confocal microscopy. The kinetic was repeated three times and representative images are shown. Scale bar: 100μm. B and C: time course of thyroid hormone receptor *CrTR* mRNA expression between 0 to 28hpf of *Ciona robusta* development (semi quantitative PCR). The extent of *CrTR* expression was compared with *CrS26* mRNA. B: a representative gel agarose image; C: the corresponding histogram is the mean of three independent experiments with standard deviation. The value set to 1 was chosen as the mean value at 18 hpf. The * represents the *P* value between 18hpf and the other time points with *P*<0.05.

### 3.2. *T4 is necessary for the initiation of the apoptotic wave and the tail regression, and modulates* CrTR *and* CrRXR *expression*

The role of T4 in the apoptotic wave was investigated by TU treatment experiments. Larvae were raised in medium treated with TU at hatching and collected either after 6h or after 12h of treatment (middle of apoptotic wave *versus* end of the tail regression period, respectively). In control larvae, the presence of TUNEL-positive nuclei is observed in 80% of larval tails at 6hph and complete tail resorption is observed in 70% of larvae at 12hph, with the remaining 30% still swimming. In TU treated larvae, neither apoptotic nuclei nor caudal regression are observed and 100% of the treated larvae are still swimming (entire tail) after 12h of treatment (Figure 2). The suppression of the apoptotic wave in *Ciona robusta* tail induced by TU is partially reversed by a simultaneous treatment with T4: after 6h of co-treatment, about 50% of larvae show a rescued phenotype with apoptotic nuclei in their tail at 6hph and go through tail regression 12h after co-treatment (Figure 2).

**Figure 2:**
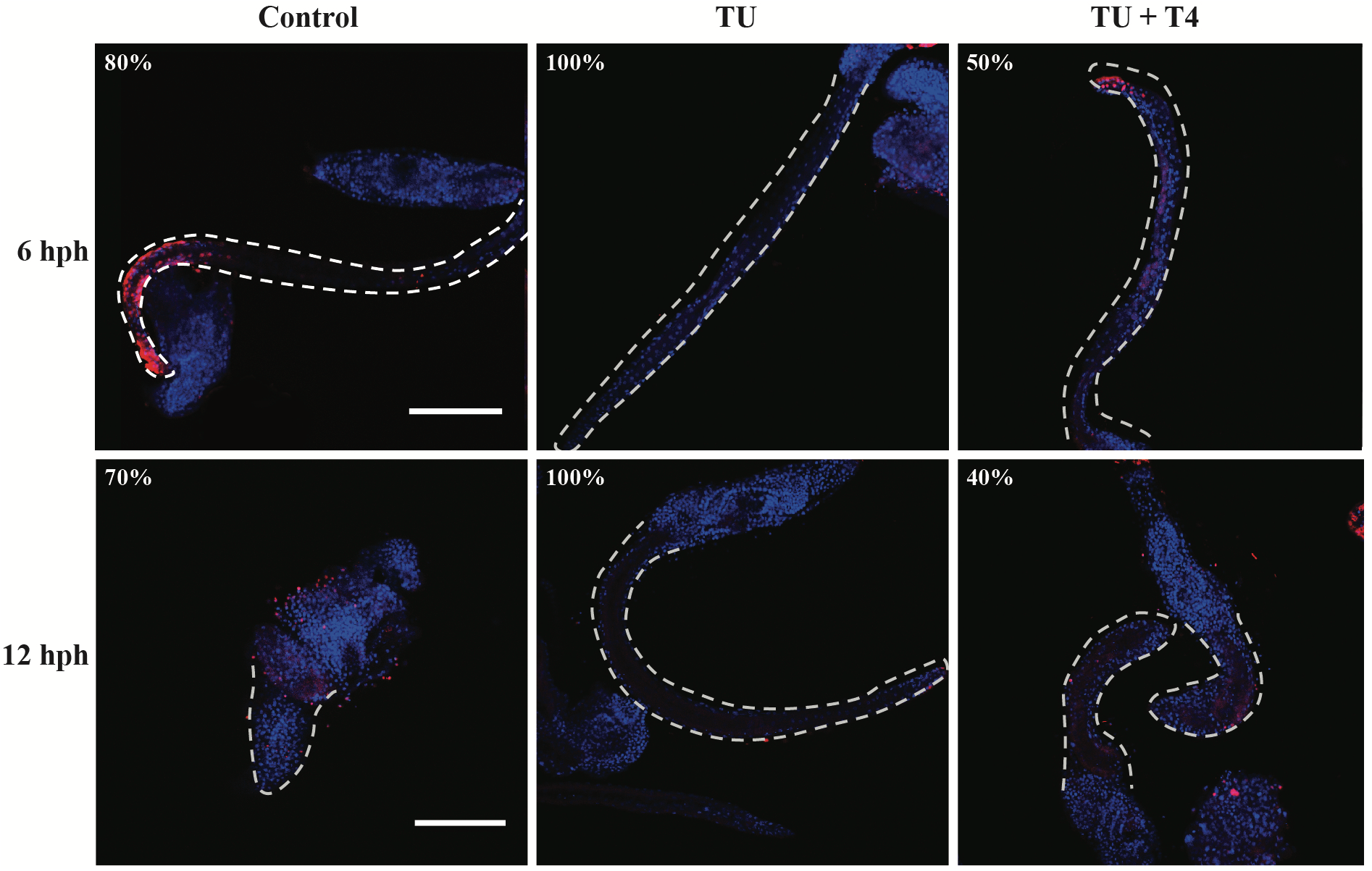
Inhibition of metamorphosis by thiourea (TU) and rescue with T4 treatment. Hatching larvae were treated (TU) or not (control) with TU 500μM and collected after 6h and 12h of treatment. Rescue larvae (TU+T4) were treated at hatching time with TU 500μM/T4 100nM mixture and collected at 6h and 12h of treatment. Fixed larvae were double-labeled with TUNEL (red) and Dapi (blue) and observed by confocal microscopy. The double labeling is shown with the tail contour drawn in grey. The experiment was repeated three times and percentage of larvae showing the presented expression pattern is indicated on each panel. Scale bar: 100μm except for 12hph Control (75μm).

To test cell-autonomy in T4-responsive tail cells, and given the progressive posterior-to-anterior initiation of apoptosis, we performed microsurgery to remove the most posterior zone of the tail. The severed larvae completely stop their metamorphosis, and show no TUNEL staining even at 12hph (Figure 3, A and B in comparison with control larva in figure 2). Tail cells therefore do not answer in a cell-autonomous way to T4 signaling. When the severed larvae are treated with T4 directly after the microsurgery, the polarized phenomenon of apoptosis is re-initiated at the posterior pole after 6h of treatment, and this metamorphosis process is re-established in 30% of severed larvae at 12hph (Figure 3A). This suggests that a signaling center is generated under T4 signaling pathway at the posterior tip of metamorphic larvae.

**Figure 3:**
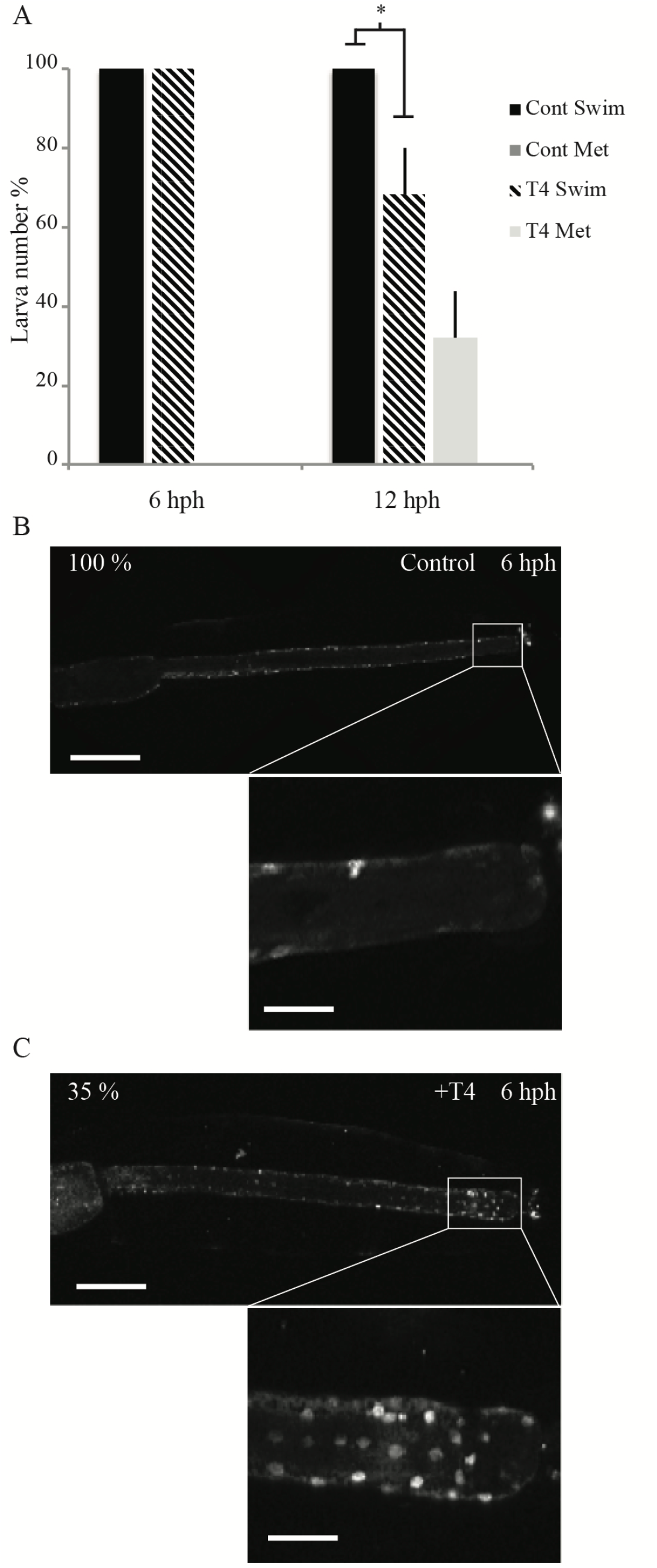
Metamorphosis of amputated larvae treated with T4. At hatching, the posterior extremity of larva tails (a quarter) was removed, amputated larvae were treated or not with T4 100nM and collected at 6h and 12h of treatment. For each kinetic point, state of 50 larvae was counted (metamorphosed *versus* swimming larvae), then collected and fixed for apoptosis detection (TUNEL, white nuclei). A: The histograms are the mean of three independent experiments with standard deviation. The * represents the *P* value between untreated and T4 treated larvae with *P*<0.05. Cont Swim: swimming untreated larvae; Cont Met: metamorphosed untreated larvae; T4 Swim: swimming T4 treated larvae; T4 Met: metamorphosed T4 treated larvae. B-C: Representative images for apoptosis detection in untreated (B) and T4 treated (C) amputated larvae are shown at 12 and 6hph, respectively. Percentage of larvae showing the presented expression pattern is indicated on each panel. Scale bar: 100μm (insert scale bar: 30μm)

Hatching larvae were treated with TU and tails were collected at different time points from hatching to the end of tail regression for qPCR experiments. *CrTR* expression decreases over time in control tails (> 7-fold decrease), but treatment with TU significantly prolongs and amplifies its expression (Figure 4A). In contrast, after hatching, the expression of *CrRXR* mRNA increases significantly until reaching a maximum at 6hph and lightly decreases at later stages (Figure 4A and Table 2). The increase of *CrRXR* in the tails is slowed down in TU-treated larvae with a significant difference at 6hph compared to the tails of control larvae (Figure 4B and Table 2).

**Figure 4:**
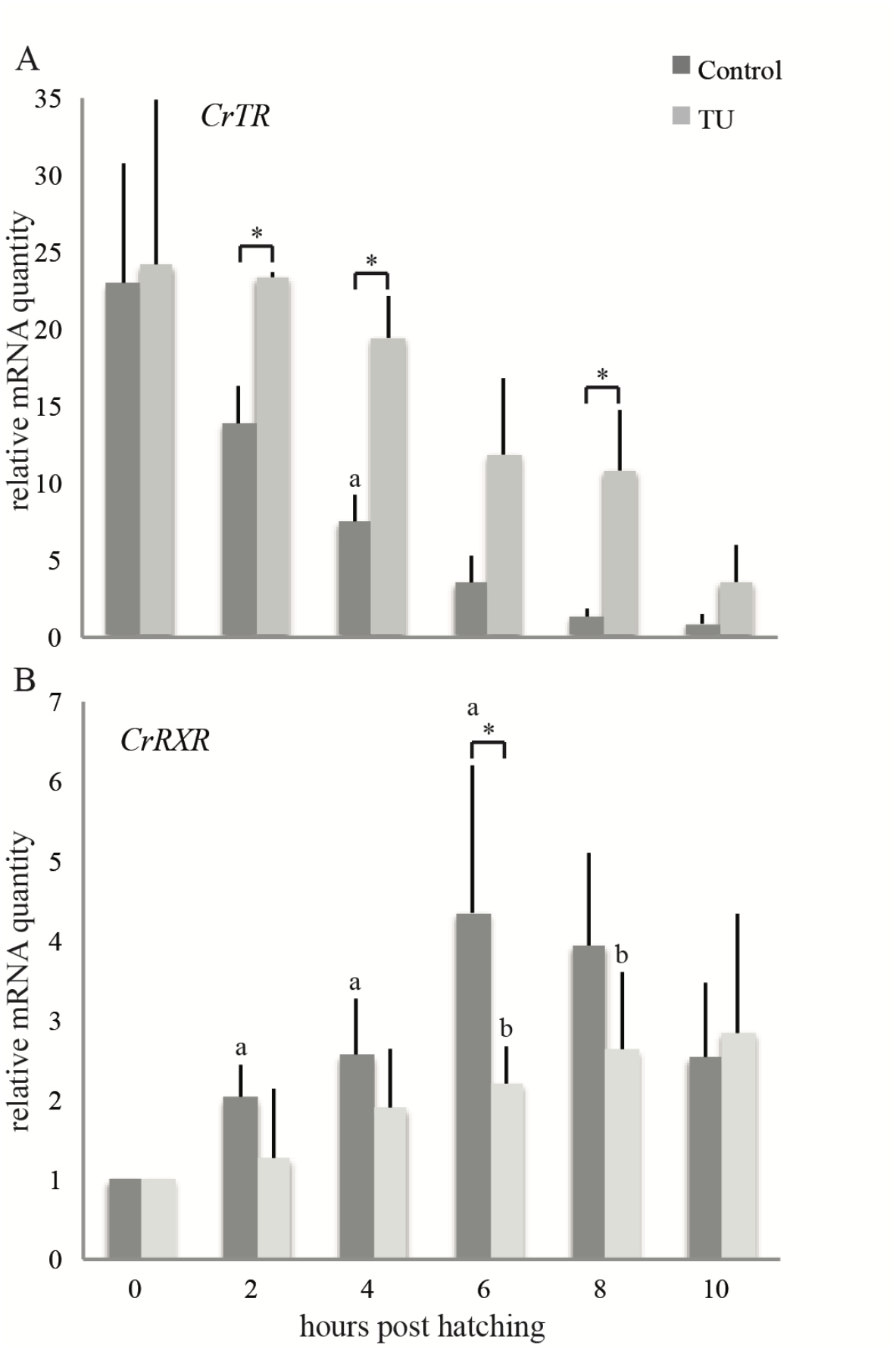
mRNA expression of *CrTR* and *CrRXR* during metamorphosis. Time course of *CrTR* (A) and *CrRXR* (B) mRNA expression from hatching to 10hph in control (dark grey) and TU treated (light grey) tails by qPCR. The histograms are the mean of three independent experiments (*i.e*. 3 different spawns with each time point corresponding to a pool of larvae) with standard deviation; data were normalized to respective *CrS26* mRNA expression values. A relative mRNA quantity value of one corresponds to the lowest amount of control target mRNA. The * represents the *P* value between control and TU treated larvae at the same stage with P<0.05. The a and b represent the *P* value between hatching and the different stages in control and TU treated larvae, respectively, with *P*<0.05.

### 3.3. *Caspase 8/9 but not Bax nor Bcl-XL expression is modulated by T4 signaling during tail regression in* Ciona robusta

Apoptotic mechanisms in vertebrates can be initiated by two classical pathways, the intrinsic (initiated by caspase-9) and extrinsic (initiated by caspases 8 and 10) pathways (Ichim & Tait, 2016). Numerous studies in amphibian metamorphosis showed that the caudal regression is under the control of the intrinsic pathway of apoptosis (Sachs *et al*., 2004; Rowe *et al*., 2005; Hanada *et al*., 2013).

In *Ciona robusta*, no caspase was specifically linked to the intrinsic or extrinsic pathway (Chambon *et al*., 2002; Dehal *et al*., 2002; Terajima *et al*., 2003; Weill *et al*., 2005) but one shows protein domains similar to both caspase-8 and -9 (Weill *et al*., 2005). This CrCasp8/9 displays two DED motifs in the pro-domain and the pentapeptide QACQG in the active site, similar to human Caspases 8 and 10 and shows a p20/p10 domain more similar to that of human Caspase 9 (Figure 5A and Weill et al., 2005). The expression of *CrCasp8/9* increases from hatching to 6hph where it reaches a peak (two-fold increase compared to at hatching) (Figure 5C). The TU treatment abolishes the increase of *CrCasp8/9* expression (Figure 5C and Table 2). In addition, expression of two key *Bcl2* family (members of the apoptosis intrinsic pathway), *Bax* and *Bcl-XL*, was detected. The expression of *CrBax* mRNA does not change over time, both in control and TU-treated larvae (Figure 5C and Table 2). The expression of *CrBcl-XL* starts to increase 6h post hatching and shows a 5-fold increase 4h later compared to hatching stage but the TU treatment does not alter significantly its expression level (Figure 5C and Table 2). In this context, the molecular mechanism implicated in the tail regression in *Ciona robusta* was investigated by first using specific functional inhibitors of caspases, *i.e*. inhibitors of vertebrate caspase-8 and caspase-9 (IETD-fmk and LEHD-fmk respectively). After 10h of treatment, only 35% of the caspase-9 inhibitor-treated larvae have strong or total tail regression while 70% of untreated larvae no longer have tails (Figure 5B). In contrast, the inhibitor of caspase-8 had no significant influence on tail regression compared to control larvae (Figure 5B).

**Figure 5:**
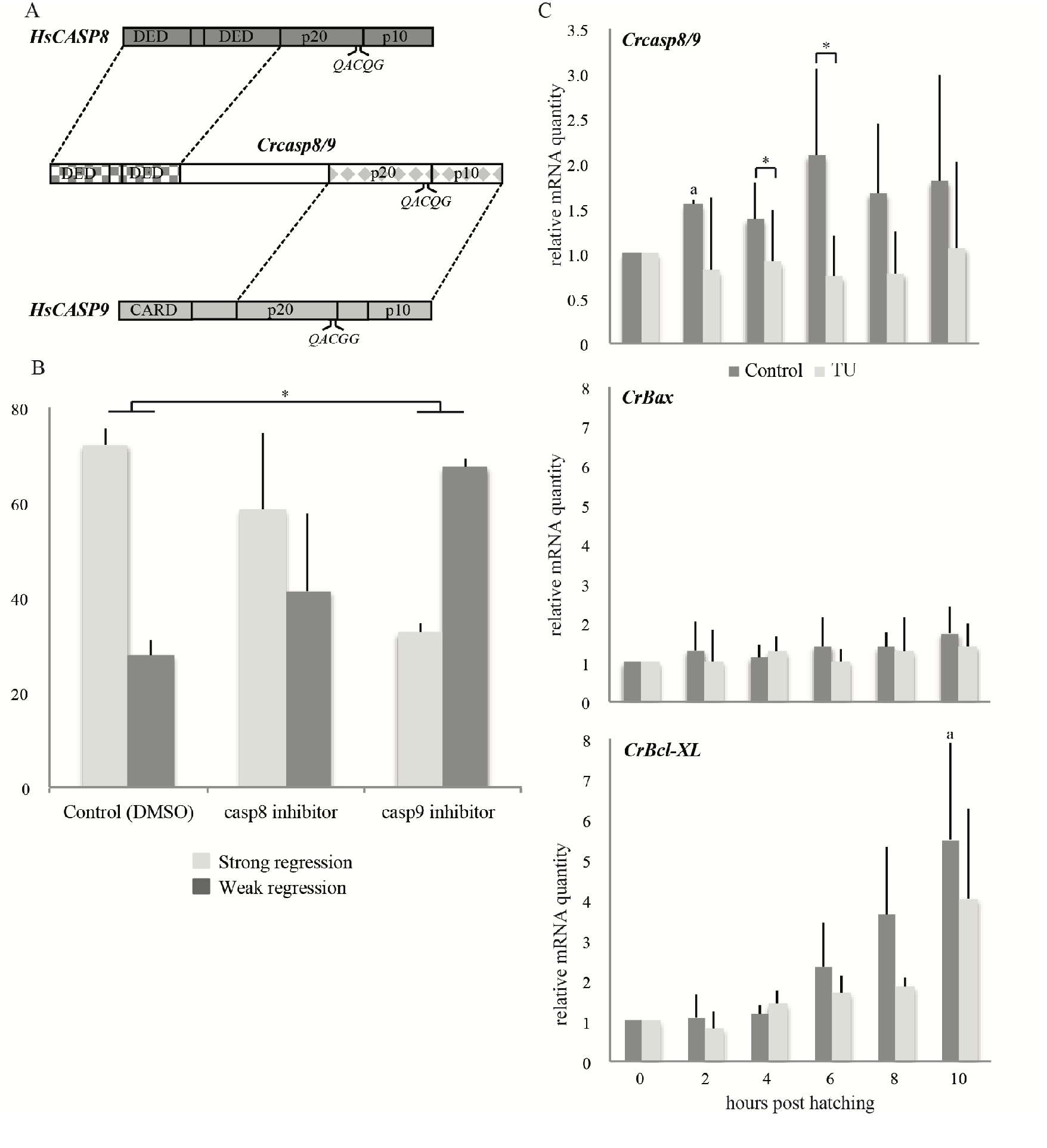
Detection of intrinsic apoptosis during metamorphosis of *Ciona robusta*. A: Schematic representation of CrCasp8/9 compared with human caspases -8 and -9 by sequence homology. B: Activity of specific caspase inhibitors. At hatching, larvae were treated with control (DMSO), 100 µM IETD-fmk (caspase-8 inhibitor) or 100 µM LEHD-fmk (caspase-9 inhibitor). Tail regression was evaluated 10h after treatment. “Strong regression” corresponds to a regression of 50 % of the tail or more (light grey) and “Weak regression” corresponds to a decrease of less than 50% of the tail (dark grey). The histograms are the mean of two independent experiments; for each treatment, state of 100 larvae was counted. C: Time course of mRNA expression of *CrCasp8/9, CrBax* and *CrBcl-XL* from hatching to metamorphosis (hph) in control (dark grey) and TU treated (light grey) tails. The histograms are the mean of three independent experiments (*i.e*. 3 different spawns with each time point corresponding to a pool of larvae) with standard deviation; data were normalized to respective *CrS26* mRNA expression values. A relative mRNA quantity value of one corresponds to the amount of target mRNA at hatching time. The * represents the *P* value with *P*<0.05. The a represents the *P* value between hatching and the different stages in control with *P*<0.05.

## 4. Discussion

### 4.1. *Thyroid hormone signaling-dependent apoptosis underlies tail regression in* Ciona robusta

In amphibians, metamorphic changes involving structural, physiological, biochemical and behavioral transformations are primarily controlled by thyroid hormone signaling and these processes can last for a few several days (Sachs *et al*., 2000). In ascidians, this process is much faster: Matsunobu and Sasakura have timed the different steps of metamorphosis in *Ciona robusta*, which from hatching to complete tail regression take only 12 hours (Matsunobu & Sasakura, 2015).

After hatching, the progression of an apoptotic wave goes from the posterior to the anterior zone of the *Ciona robusta* tail. Here, we show that the inhibition of T4 signaling by TU blocks the initiation of the apoptotic wave and subsequent tail regression, suggesting that T4 is necessary to initiate apoptosis (Figure 2). This T4 effect, consistent with that found in amphibians (Sachs *et al*., 1997a; b), is confirmed by the results of our rescue experiments where apoptosis and subsequent tail regression is reactivated by T4 co-treatment with TU (Figure 2).

We show that T4 acts in the initiation of apoptosis *via* activation of an intermediate signaling center located in the tail end. This is congruent with reports of a number of genes being only expressed at the posterior end of the tail of *Ciona robusta* such as *Sccpb* (similar to selectin P), a gene under the MAPK signaling pathway (Cr-ERK and Cr-JNK) that is essential for apoptosis-dependent tail regression (Chambon *et al*., 2007). Interestingly, ERK is activated by phosphorylation only at the tail end before the wave of apoptosis (Chambon *et al*., 2002; Krasovec *et al*., 2019). As a consequence, we postulate that the posterior part of the larval tail contains the signaling source responsible for the initiation of observed wave of apoptosis and the metamorphosis process, and that this signaling center is under the control of T4 signaling. In our tail removal experiments (Figure 3), T4 treatment probably allows local and fast expression/recruitment of proteins specific to the end of tail and then apoptosis recovery at 6hph (Figure 3). Thyroid hormone signaling therefore appears to activate the apoptosis pathway in a directional way in the period preceding adhesion by initiation of the posterior signaling center. The link between thyroid hormone signaling and apoptosis is further documented by the results of qPCR experiments showing a thyroid hormone-dependent transcriptional induction of *CrCasp8/9* during tail regression (Figure 5C).

### 4.2. *Apoptosis during* Ciona robusta *tail regression does not rely on a classical intrinsic pathway*

Thyroid hormone-dependent apoptosis in *Xenopus* tadpole involves the intrinsic pathway (Sachs *et al*., 1997a; b; Xiong *et al*., 2014; Ichim & Tait, 2016). Our caspase-inhibition treatments support a caspase-9-like activity in the initiation of apoptosis, therefore similar to the function of the intrinsic pathway. However, this experiment cannot identify the exact active caspase involved in the process since the active site of the CrCasp8/9 is comparable to that of human caspase-8 (Figure 5A). Our qPCR results show an important induction of *CrBcl-XL* starting at 6hph, when *CrCasp8/9* mRNA expression is maximal. This result is surprising because the vertebrate Bcl-XL is known for its anti-apoptotic action (for review see (Cui & Placzek, 2018). If this activity is conserved in chordates, the induction of *CrBcl-XL* might have a function in the conservation of an anti-apoptotic state for the cells that migrate to the trunk during tail regression. This is in contrast to the *Xenopus* metamorphosis where Bcl-XL has no apparent function in the thyroid hormone-induced apoptosis (Johnston *et al*., 2005) and where cell migration from the tail to the body has never been reported (see review Yaoita, 2019).

### 4.3. *New model of interaction between thyroid hormone and apoptotic pathway in* Ciona robusta

From our results, the interaction between thyroid hormone signaling and the apoptosis in the tail of *Ciona robusta* appears different from that described in *Xenopus* tadpoles. A concomitance between high levels of T4, CrTR and apoptosis is not observed in *Ciona robusta* in contrast to *Xenopus laevis* (Figure 6 according to the work of (Sachs *et al*., 2000; Ishizuya-Oka *et al*., 2010)). Our results indicate that *CrTR* expression is high at hatching, when *CrRXR* expression and apoptosis are still low or not detected, respectively. After hatching, thyroid hormone and *CrTR* levels progressively decrease concomitantly with increasing quantities of apoptotic nuclei, *CrRXR* and *CrCasp8/9* mRNAs (Figure 6). These results suggest that CrTR behaves like TRα in amphibians where, in the absence of THs, it prevents the progression of metamorphosis and promotes the growth of tadpoles (Wen & Shi, 2015). Transcription of both *CrTR* and *CrRXR* genes is modified by the treatment with TU, arguing for thyroid hormone-dependent expression of both receptors.

**Figure 6:**
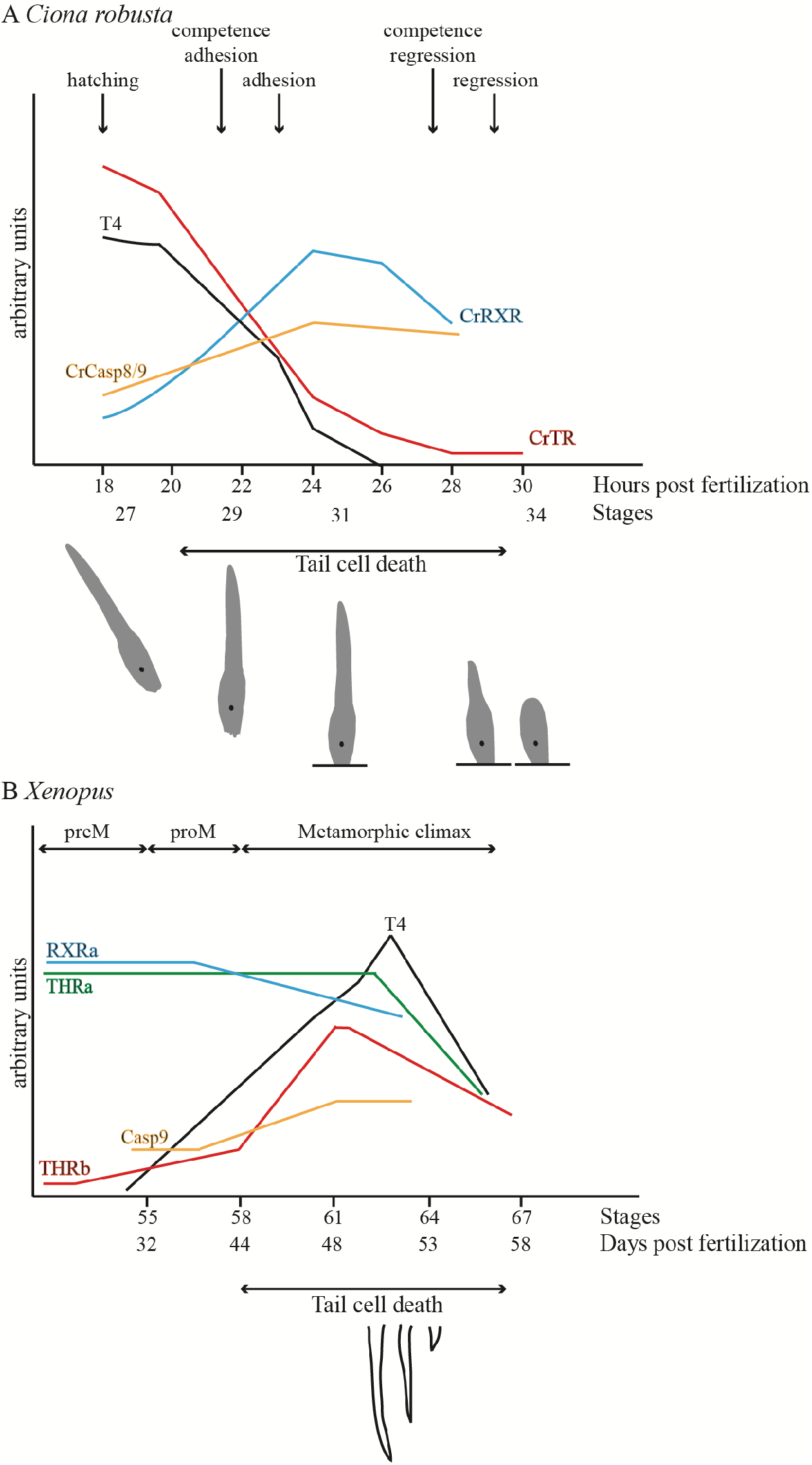
Comparative scheme between *Xenopus* and *Ciona robusta* tadpole metamorphoses. Summary of developmental stage-dependent expression of TR and RXR genes and tail cell death of *Ciona robusta* tadpole (A) in comparison to *Xenopus* tadpole (B) during metamorphosis. preM and proM, pre- and prometamorphosis, respectively (adapted from (Sachs *et al*., 2000; Rowe *et al*., 2005; Ishizuya-Oka *et al*., 2010; Matsunobu & Sasakura, 2015; Hotta *et al*., 2020).

However, the results of these qPCR experiments do not allow to conclude on the functionality of the TR/RXR heterodimer which will require further *in vitro* study of nuclear receptor activities.

### 4.4. Conservation of thyroid hormone and metamorphosis in chordates

Metamorphosis is an ancestral feature of chordates with conserved molecular determinism (Holzer *et al*., 2017). Thyroid hormones and their receptors have been systematically involved in metamorphosis in all chordates in which their role has been tested, even in animals located at very distant branches of the chordate phylogenetic tree (amphibians, teleost fishes and amphioxus). Different larva-to-juvenile transitions described can be considered as homologous, based on the conservation of the thyroid hormone signaling pathway generated by thyroid hormones and their receptors. We have shown in this study that the thyroid-dependent metamorphosis of *Ciona robusta* has a different mode of operation from what is classically known. Interestingly, another organism also stands out from the other chordates: in the sea lamprey, it is also the drop of circulating T4 that enables its metamorphosis, which shows features of apoptosis in the epithelial cells of the biliary tract (Boomer *et al*., 2010; Morii *et al*., 2010) and the pronephric kidney (Ellis & Youson, 1990). But in contrast to *Ciona*, exogenous THs slow down, rather than accelerate, the natural lamprey metamorphosis with binding to their specific receptors (Youson, 1997; Manzon *et al*., 2014; Manzon & Manzon, 2017) (Figure 6). Despite being a more distant vertebrate relative compared to urochordates (Delsuc *et al*., 2006), the amphioxus has a functional thyroid hormone receptor, and THs induce metamorphosis in amphioxus as in the frog, despite apoptosis was not shown during the metamorphic process (Paris *et al*., 2008). In connection with these comparisons, it is important to note that metamorphosis in *Ciona* lasts only a few hours (instead of several days for the amphioxus, lamprey or xenopus) and in this regard, is more comparable with that of echinoderms within the deuterostomes.

### 4.5. Conservation of thyroid hormone and metamorphosis in bilaterians

Within the deuterostomes, the sea urchin (echinoderm) transforms from a swimming larva into a sessile juvenile (bilateral symmetric larva into radial symmetric and benthonic adult) in a few hours (Sato *et al*., 2006). All eight arms of the sea urchin larva reduce from their tip to the trunk by apoptosis when treated with THs, and inhibition of apoptosis prevents the induction of metamorphosis (Saito *et al*., 1998; Lutek *et al*., 2018; Taylor & Heyland, 2018; Wynen & Heyland, 2021). Similar to *Ciona robusta*, the sea urchin has 1 RXR and 1 TR (Howard-Ashby *et al*., 2006), exogenous TH treatment accelerates its metamorphosis but transcription of target genes via its TR is not activated by THs (Taylor & Heyland, 2018). Recently, TH action in sea urchin has been shown to act via the MAPK-ERK1/2 pathway (Taylor & Heyland, 2018) as it was shown for *Ciona robusta* (Chambon *et al*., 2007). These similitudes between urochordate and echinoderm metamorphic processes suggest an ancestrally conserved regulation by thyroid hormone, through the MAPK-ERK pathway and involving TR as a constitutive transcriptional regulator, therefore differing significantly from the “classical” vertebrate thyroid hormone signaling.

Outside of deuterostomes, in both oysters and mussels, THs peak at the gastrula stage and decrease just after the trochophore stage. This variation is correlated with the presence of the TR (absent after the trochophore stage). The oyster has 1 RXR and 1 TR, and its TR inhibits its own expression supporting the transcriptional repression activity of the TR (Huang *et al*., 2015). In addition, two caspases play a key role in the loss of the foot and velum during larval metamorphosis (Yang *et al*., 2015). In mussels, while TH treatment accelerates metamorphosis, knockdown of its TR leads to inhibition of metamorphosis: thus, the TR may have transcriptional repression activity affecting competence for the metamorphic transition (Li *et al*., 2020) (figure 6). This TH-dependent signaling pathway with TR acting as a constitutive transcription factor may therefore be a probable bilaterian ancestral regulatory pathway. Both this bilaterian and the vertebrate regulatory pathways may have co-existed in the last common ancestor of chordates, and differentially selected for: the “vertebrate”-type was conserved in amphioxus and vertebrates, while the “bilaterian”-type was conserved in *Ciona* (Morthorst *et al*., 2022).

## 5. Conclusions

Collectively, these results suggest that thyroid hormones are involved in the initiation of the apoptotic wave leading to tail regression in *Ciona robusta*, similar to previous findings in amphibians. However, in *Ciona robusta* the mode of action shows great variation compared to the classically described scheme in vertebrates, both at the level of the thyroid hormone/TR interactions and also at the level of the apoptotic pathway. Despite a phylogenetic position in the vertebrate sister group, *Ciona robusta* has retained the ancestral thyroid hormone pathway, i.e. a non TH/TR interaction but with a constitutive TR that represses progression to metamorphosis by promoting tadpole growth. This significance of the conservation but also these differences might be linked to the evolution of a very rapid metamorphosis in an organism of simple architecture.

## Abbreviations

T3: tri-iodothyronine
T4: thyroxin
TR: thyroid hormone receptor
TU: thiourea
hpf: hours post-fertilization
hph: hours post-hatching.

## Acknowledgments

We are very grateful to Philippe Clair for his expertise on qPCR experiments, Vicky Diakou for her expertise on confocal microscopy and Julien Claude for his help on statistical analyses. Some data used in this study were produced using the Montpellier RIO imaging platform (Confocal microscopy) and the qPHD UM2/GenomiX platform (qPCR experiments) (Montpellier, France). We acknowledge the imaging facility MRI, member of the national infrastructure France-BioImaging supported by the French National Research Agency (ANR-10-INBS-04, «Investments for the future»).

## Legends of figures

**Supplementary data:**
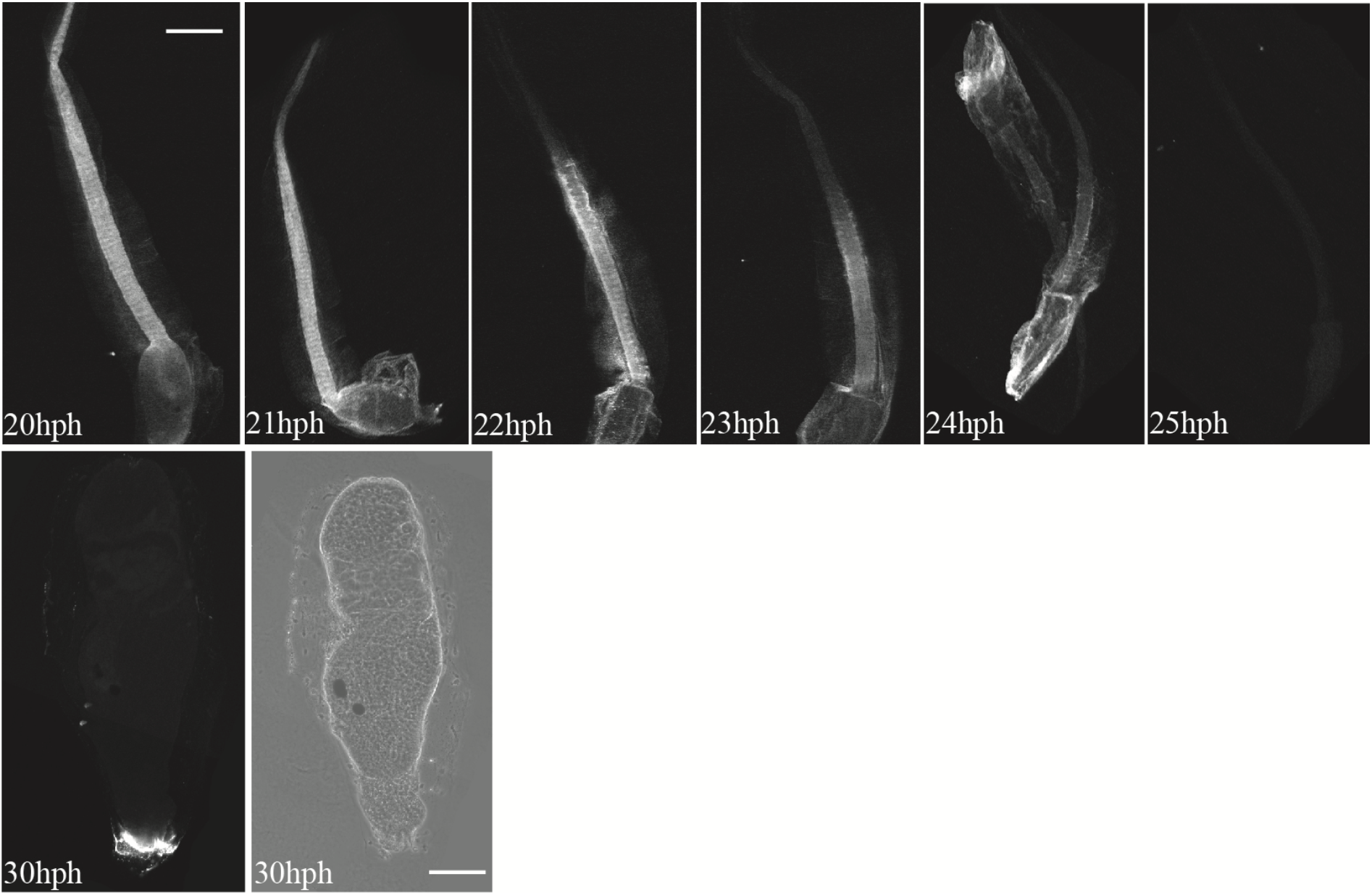
T4 staining (white) at different stages of development, from hatching to the end of tail regression (hph: hour post-hatching) by confocal microscopy. The kinetic was repeated three times and representative images are shown. Scale bar: 100μm except for late tail regression at 30hph (25μm).

